# Cutting across structural and transcriptomic scales translates time across the lifespan and resolves frontal cortex development in human evolution

**DOI:** 10.1101/2020.08.06.240077

**Authors:** Christine J. Charvet

**Author notes:** Corresponding author Christine Charvet, PhD, Center for Neuroscience, Delaware State University, Dover, DE, 19901, USA.

## Abstract

How the unique capacities of human cognition arose in evolution is a question of enduring interest. It is still unclear which developmental programs are responsible for the emergence of the human brain. The inability to determine corresponding ages between humans and apes has hampered progress in detecting developmental programs leading to the emergence of the human brain. I harness temporal variation in anatomical, behavioral, and transcriptional variation to determine corresponding ages from fetal to postnatal development and aging, between humans and chimpanzees. This multi-dimensional approach results in 137 corresponding time points across the lifespan, from embryonic day 44 to ∼55 years of age, in humans and their equivalent ages in chimpanzees. I used these data to test whether developmental programs, such as the timeline of prefrontal cortex maturation, previously claimed to differ between humans and chimpanzees, do so once variation in developmental schedules is controlled for. I compared the maturation of frontal cortex projections from structural magnetic resonance (MR) scans and from temporal variation in the expression of genes used to track long-range projecting neurons (i.e., supragranular-enirhced genes) in chimpanzees and humans. Contrary to what has been suggested, the timetable of prefrontal cortex maturation is not unusually extended in humans. This dataset, which is the largest with which to determine corresponding ages across humans and chimpanzees, provides a rigorous approach to control for variation in developmental schedules and to identify developmental programs responsible for unique features of the human brain.

## Introduction

Humans differ in many respects from other primates, but the developmental programs that gave rise to features unique to the human brain are still unclear. Debates have centered on whether humans possess an extended duration of prefrontal cortex white matter growth, myelination, and synaptogenesis relative to chimpanzees and other primates [1–8]. These controversies have emerged in part because there is no systematic approach to compare ages across the lifespan in humans and chimpanzees. Developmental schedules vary widely across species, with overall human developmental schedules extended for an unusually long time compared to those of many other mammals and possibly chimpanzees [1,8–14].

Controlling for variation in developmental schedules is required to determine whether the unfolding of developmental programs, such as cortical projection or myelination, occurs for an unusually long time in humans relative to chimpanzees [8, 14]. The lack of a systematic approach to compare ages across the lifespan of primates has hampered progress in our ability to identify developmental programs that have evolved within the human lineage. To overcome these limitations, I developed a multi-scale approach with which to determine corresponding ages across species. I harness temporal variation in transcriptional, anatomical, and behavioral variation to identify corresponding ages across the lifespan in humans and chimpanzees. The present study builds on a previously developed line of work (i.e., translatingtime.net), which considers the timing of developmental transformations (i.e., abrupt changes in developmental processes) to identify corresponding ages between model organisms and humans [8, 14–20]. The translating time project relied on abrupt changes that occur over the course of fetal development (cell birth, synaptogenesis, myelination) as a basis with which to find corresponding time points across humans and model organisms. While this approach has served to identify corresponding ages during fetal development (e.g., mice, macaques, cats), new approaches are needed to determine corresponding ages throughout postnatal ages, including aging. Capturing temporal variation in transcription, anatomy, and behavioral variation from dynamic time warping, non-linear regressions, and abrupt changes in biological programs ensures concordance of identified time points and finds corresponding ages across the lifespan.

A number of developmental programs have been proposed to be modified within the human lineage. These include a possible extended duration of prefrontal cortex white matter maturation, myelination, and synaptogenesis, compared with that of chimpanzees [1–7]. These studies highlight the role of protracted development in establishing connections and maturation of projections supporting neural structures unique to humans. The developmental time course of connection formation and projection maturation may simply be the product of an extended duration of development in humans relative to other species [20-24]. It is therefore not clear whether the duration of connection formation and projection maturation is extended in humans relative to chimpanzees once variation in developmental schedules is controlled for across species.

I used a two-pronged approach to compare the developmental time course of frontal cortex circuitry maturation across species. I compared the timeline of white matter growth from structural magnetic resonance (MR) scans because the white matter houses long-range projections and from temporal variation in the expression of genes used as long-range projection markers in humans and chimpanzees. I use supragranular-enriched genes, which encode filaments (e.g., *NEFH*), synapse components (e.g., *VAMP1*), and voltage-gated channels (e.g., *SCN4B*), as markers of long-range projections [25]. Temporal variation in the expression of these supragranular-enriched genes aligns with the growth of white matter pathways across species [20]. The expression of select supragranular-enriched genes evolves with modifications in tractography from diffusion MR imaging [20, 26, 27]. Finally, the expression of supragranular-enriched genes covary with functional connectivity across the human brain [28]. Together, these studies highlight the utility of supragranular-enriched genes as markers of long-range projecting neurons to test whether prefrontal cortex circuitry development is extended in humans relative to the timing of other biological pathways. Together, the integration of these scales ensures accuracy of identified time points and yields a more complete understanding of the evolution and development of frontal cortex circuitry in the human lineage [21, 27].

Surprisingly, many developmental programs are either conserved across humans and chimpanzees or occur earlier in humans than in chimpanzees [8]. Weaning stands our as occurring unusually early in humans relative to the timing of most other developmental programs [22]. I found no evidence that prefrontal cortex maturation and synaptogenesis occurs for an unusually long time in humans. This study demonstrates the utility of a multi-scale approach in identifying corresponding time points across the lifespan and provides a baseline with which to identify modified biological programs within the human lineage.

## Materials and methods

I gathered corresponding time points (n=137) from temporal changes in gene expression, anatomy, and behavior in humans and chimpanzees (Fig. 1). Some of these time points are from abrupt changes while other time points are captured from gradual changes span multiple time points. Ages were extracted from dynamic time warping applied to temporal variation in transcription (Fig. 2), and diffusion MR data (fractional, radial and medial diffusivity). In addition, I used non-linear regressions as well as abrupt transformations to extract corresponding time points between humans and chimpanzees [2,5,6,26–46; Fig. 3, Fig. S1-S5; Table S1-S5]. All statistics were performed with the software programing language R.

**Fig. 1.**
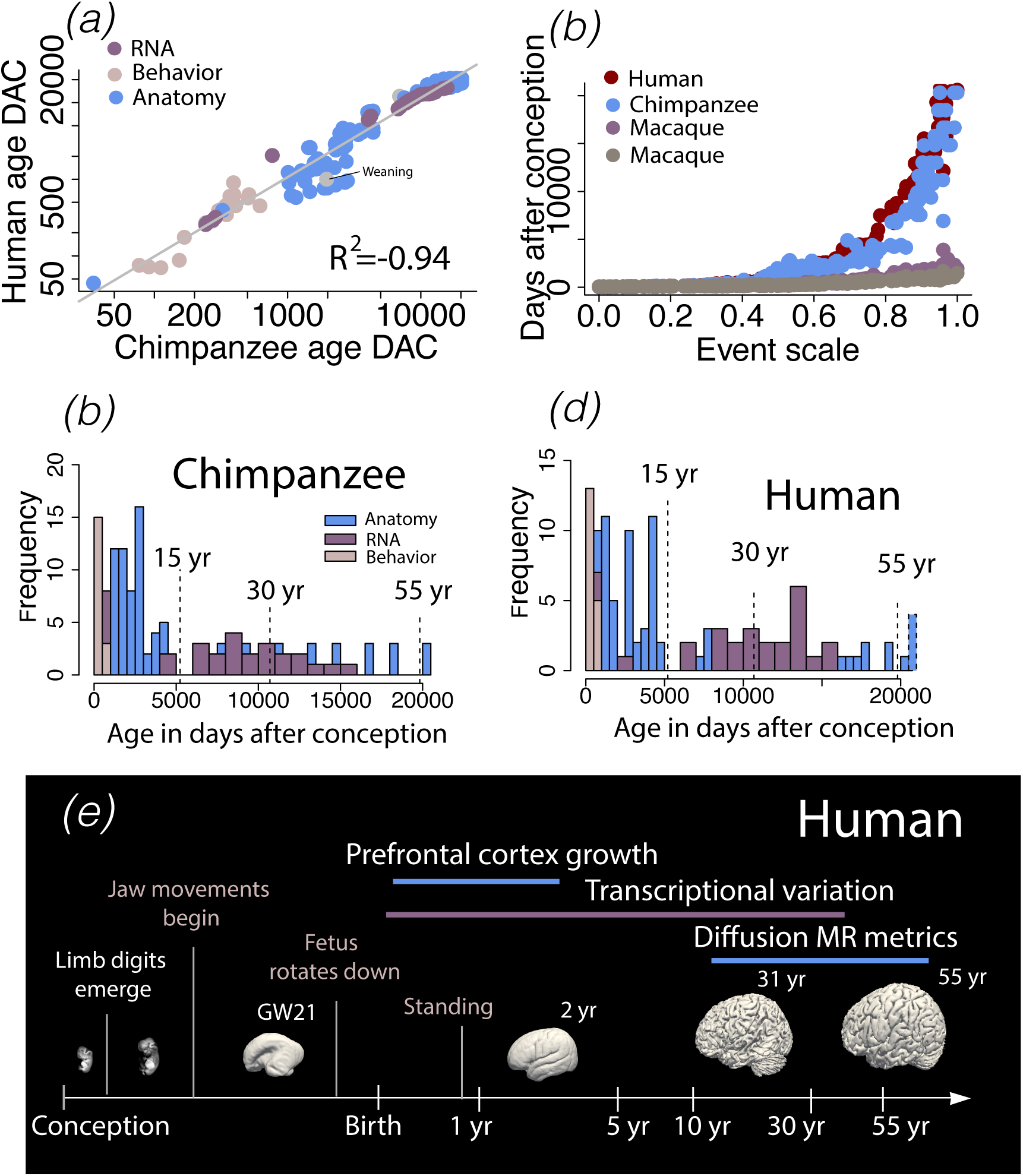
*(a)* I fit a general linear model through the natural log of age expressed in days after conception (n=137) in humans versus chimpanzees. *(b)* I also fit the natural-log-transformed values of age in days after conception versus an event scale in humans, chimpanzees, macaques, and marmosets. *(c-d)* Histograms show the age ranges in humans and chimpanzees from *(a)*. These data reveal corresponding ages of up to 55 years in humans and their corresponding ages in chimpanzees. *(e)* Some examples of biological programs and time points used to find corresponding ages are shown in humans. Surfaces were created from structural MR scans of humans at different ages.

**Fig. 2.**
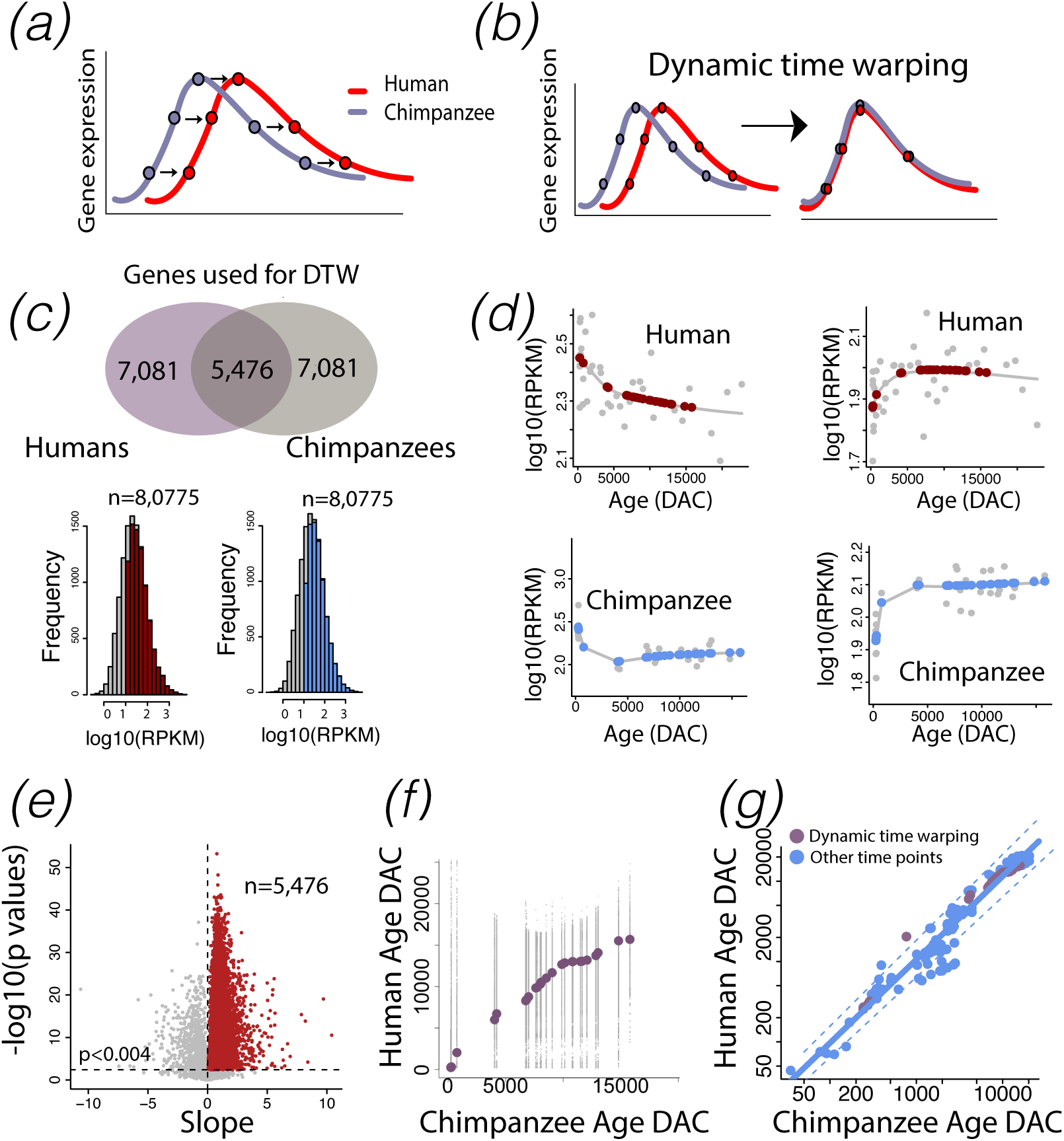
I used optimal alignments of temporal trajectories in transcription (*a-b*) to find corresponding time points across humans and chimpanzees. (*c*) Several filtering steps reduced the number of genes (n=5,476) available for dynamic time warping. I filtered genes with minimum average expression of log10(RPKM)>1 (n=8,075) per species. (*d*) I then fit a smooth spline through log-transformed RPKM+1 versus the log-transformed ages expressed in days after conception in each species to compare normalized gene expression across identical ages and time points in each species (n=31 in humans; n=31 in chimpanzees). For each gene, I fit a linear model through the log10(RPKM) of humans and age-matched chimpanzees. This approach selected genes with similar temporal variation between the two species. I corrected for multiple testing with a false discovery rate (FDR) BY threshold p value set to 0.05 to select genes that significantly covaried, which yielded 5,476 genes available for dynamic time warping. (*f*) For each gene, I applied dynamic time warping to log10(RPKM) in humans versus age-matched chimpanzees and extracted corresponding time points from these data. (*g*) I computed the median corresponding time points for all tested genes. Time points extracted from transcriptional variation (with the exception of 1 time point) fell within the 95% confidence intervals of the remaining time points, which demonstrated the concordance of identified corresponding ages from transcription to behavior and anatomy. Dashed lines indicate 95% confidence intervals generated from non-transcriptional time points in humans and chimpanzees. DAC: days after conception. DTW: dynamic time warping.

**Fig. 3.**
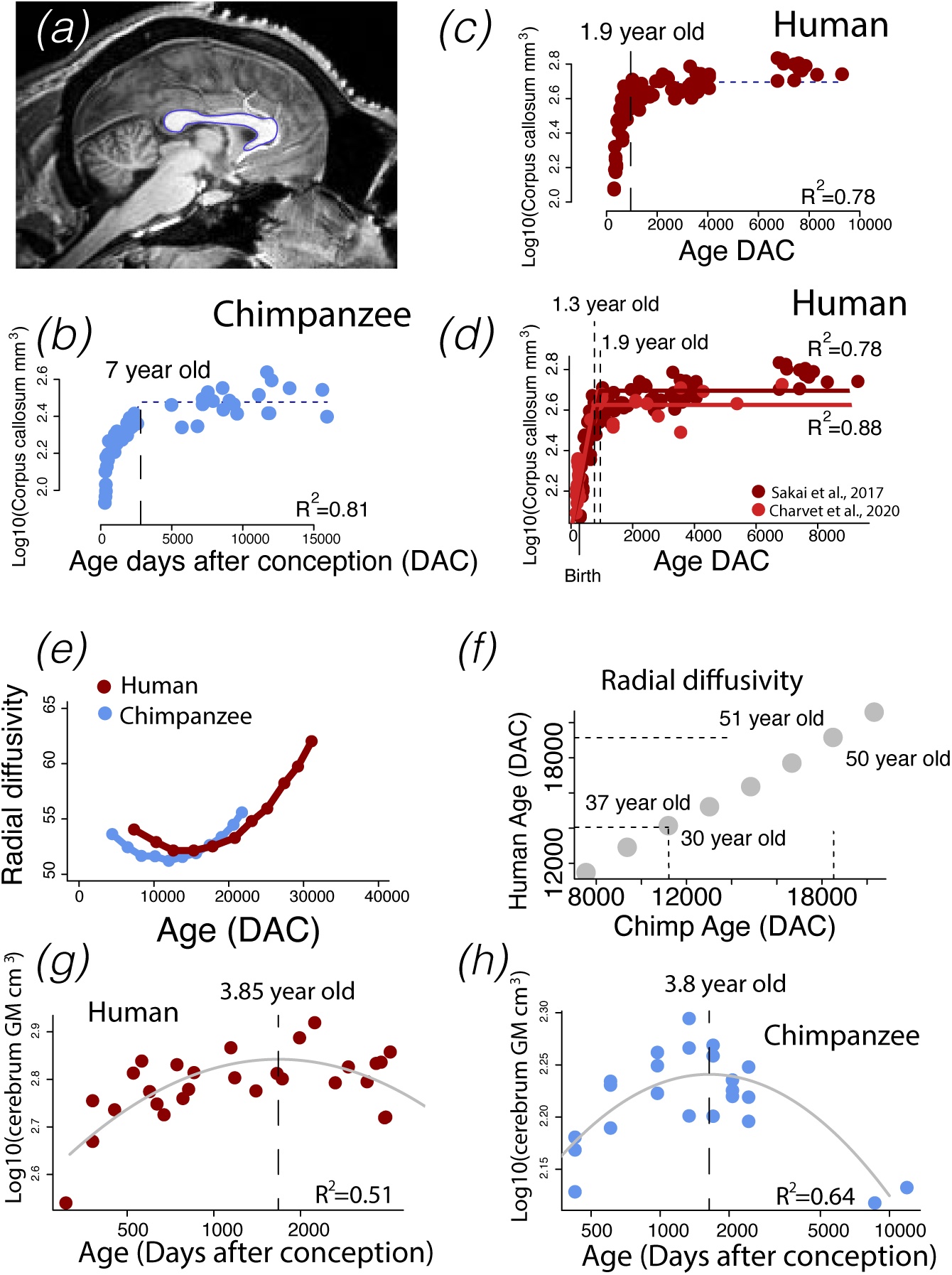
I use multiple statistical approaches to identify corresponding time points in humans and chimpanzees, a few of which are shown. I captured corresponding ages with non-linear regressions (*a-d*), dynamic time warping (*e-f*), and inflexions (i.e., peaks) in anatomical phenotypes (*g-h*). As an example, I applied nonlinear regressions to corpus callosum area measurements in humans and chimpanzees as shown from a mid-sagittal slice of a T1 structural MR chimpanzee brain scan (*a*). I identified when the corpus callosum ceases to grow in humans and chimpanzees as a corresponding time point. I also used dynamic time warping applied to variation in diffusion MR imaging (i.e., fractional anisotropy, radial diffusivity, and medial diffusivity) to find corresponding time points across species. I aligned temporal variation in radial diffusivity to find corresponding time points between humans and chimpanzees. In addition, I extracted corresponding time points from peaks in cortical gray matter volumes in humans and chimpanzees.

To compare schedules of development and aging across primates, I acquired timepoints from marmosets (n=31) and macaques (n=56; 16–18,31,34], which yeilded 195 time points across the studied primate species. Not all time points are available across the four primate species. I therefore imputed logged time points across humans, chimpanzees, macaques, and marmosets with the linear regression boot-strap method implemented with the library package mice. These imputed data served to calculate the event scale. The event scale is an ordering of developmental transformations with early events assigned a low score and late events a high score and ranges from 0 to 1. The event scale is calculated by subtracting the timing of each developmental transformation averaged across species from the earliest event and dividing these values by the difference between the latest event and the earliest event [19]. I then fit a linear model with the log10 days after conception versus the event scale with species as a factor and an interaction between species and the log10 days after conception.

### Temporal variation in transcription to determine corresponding time points

I captured corresponding time points from temporal variation in transcription in humans and chimpanzees (Fig. 2*b*). Tissue was extracted from the prefrontal cortex of humans (n=38, ∼birth to ∼61.5 year old) and chimpanzees (n=31, birth to ∼42.5 years old; [45]). Sequence reads were mapped to reference genomes hg19 and panTro3. A principal component analysis of the log-transformed reads per kilobase per million (RPKM) across the two species revealed no outliers (Fig. S1; Table S5). I selected genes with a minimum average log10(RPKM)>1 across both species, which yielded 8,075 expressed genes available for downstream analyses (Fig. 2*c*). Although the age ranges in humans and chimpanzees overlapped, they were not identical. To extract RPKM values at equivalent ages for both species, I fit a smooth spline through the log-transformed RPKM versus the log-transformed ages expressed in days after conception in chimpanzees and in humans (Fig. 2*d*). I extrapolated log10 RPKM values in humans to match corresponding time points in chimpanzees (n=31 human; 31 chimpanzees). I also extracted predicted values from smooth splines in chimpanzees so that the variance in normalized gene expression would be similar across both species (Fig. 2*d*). I fit a linear model through the log-transformed RPKM values in humans and chimpanzees iteratively for each gene. This was to select genes with significant temporal patterns in expression across the two species. I corrected for multiple testing with a false discovery rate (FDR) Benjamini & Yekutieli (BY) threshold p value set to 0.05, which yielded 5,476 genes for downstream analyses (Fig. 2*e*). I use the library dtw to warp the log-transformed RPKM values across age-matched humans and chimpanzees to find optimal corresponding time points for each gene (Fig2.*e*). I then computed the median across these data, which I use as corresponding time points in humans and chimpanzees (Fig. 2f*-g*).

### Temporal variation in diffusivity to determine corresponding time points

I applied dynamic time warping to fractional anisotropy (FA), medial diffusivity (MD), and radial diffusivity (RD) to find corresponding ages across humans and chimpanzees (Fig. S5). Identical and equidistant ages were selected from smooth splines fit to FA, RD, and MD versus age expressed in days after conception in humans and chimpanzees. I mapped these time series data to find the optimal alignment of ages across humans and chimpanzees (Fig. 1, S5), which yielded a total of 24 time points across the two species.

### Non-linear regressions to find corresponding time points

I used nonlinear regressions to capture when various brain regions cease to grow to find corresponding ages across species. Some of these data were extracted from previous studies [2, 41-42, 46] and from structural MR scans of human and chimpanzee brains. I measured the cross-sectional area from mid-sagittal measurements of the corpus callosum and the prefrontal cortex white matter from T1 MR scans of chimpanzee brains with ImageJ software (Tables S2-S3). I binarized the images to measure bilateral prefrontal cortex white matter volumes anterior to the corpus callosum in accord with previous work [2, 6]. The chimpanzee MR scans were from the National Chimpanzee Brain Resource, and the human MR scans were from [47]. I used a linear or quadratic plateau implemented with the package easynls (model=3 or 4) to capture the age of brain region growth cessation in each species. To ensure that the ages of growth cessation were not driven by outliers, I randomly subsampled individuals and extracted the age of growth cessation from these data (Fig S2-S4). I also ensured concordance of corresponding time points across data sets (Fig. 2, Fig. S4).

### Testing for protracted PFC maturation in humans with supragranular-enriched genes

In addition to testing for deviations in the timetable of white matter pathway maturation across species, I compared temporal trajectories in transcription from markers of long-range projections [39]. I considered supragranular-enriched genes, which encode filaments (e.g., *NEFH*), synapse components (e.g., *VAMP1*), and voltage-gated channels (e.g., *SCN4B*; 25-28]. I compared coefficients of variation between the two species and at multiple age ranges [20]. Comparing coefficients of variation (i.e., ratio of the standard deviation to the mean) assumes that genes that vary more in their expression in one species relative to another at late stages of development reflect protracted developmental programs of long-range projections (Fig. 4). I fit a smooth spline through the log10 of FPKM versus the log10 of age expressed in days after conception for each gene and for each species. I added a value of 1 to all log10-transformed FPKM values to retain genes that were not expressed at a particular time point. I computed the coefficient of variation for each gene across all age ranges or between 10 to 50 years of age for each species. I then performed a t-test on the coefficient of variation to identify whether variation in supragranular-enriched gene expression is significantly increased at late stages of development in humans compared with chimpanzees.

**Fig. 4.**
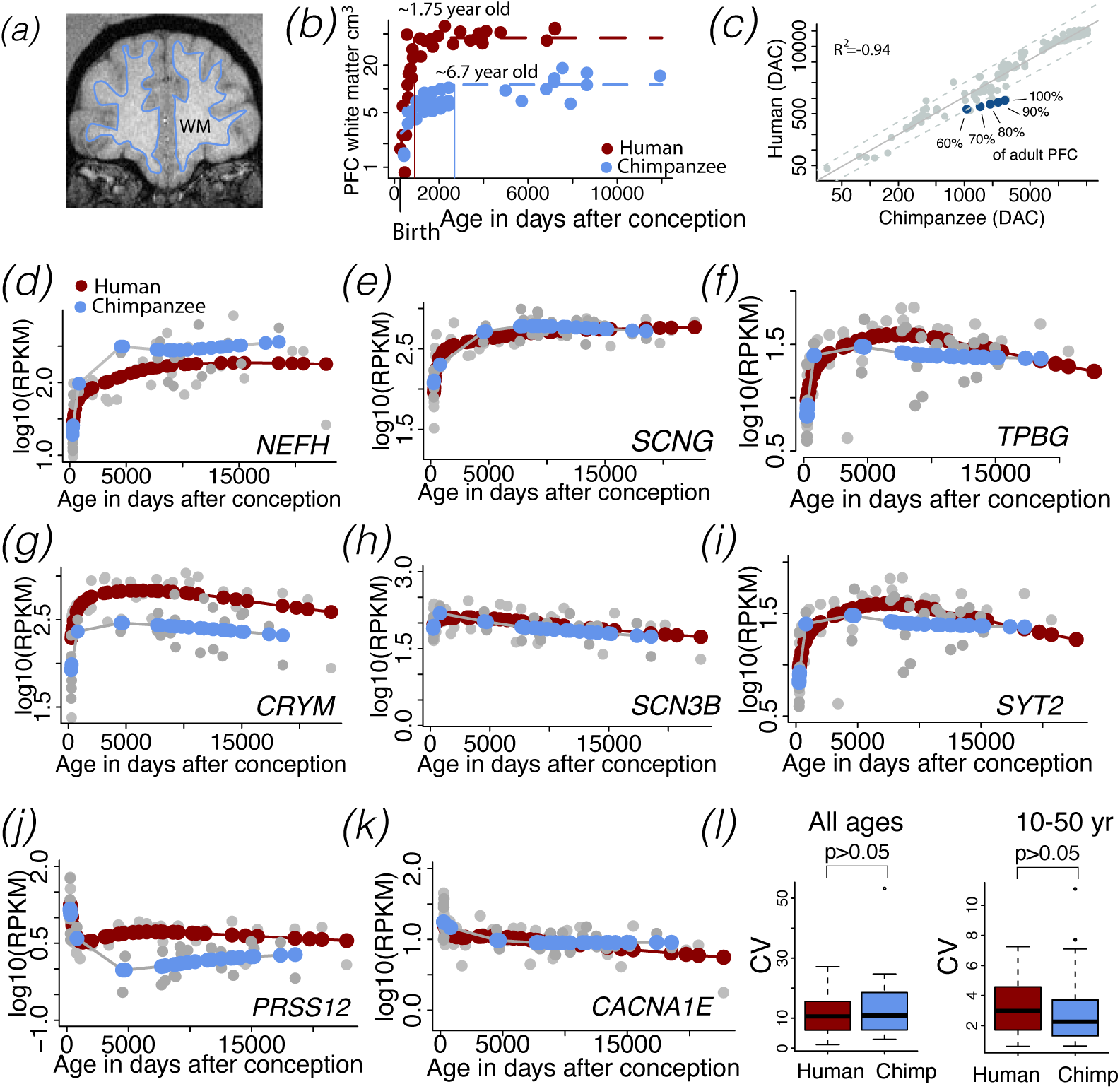
The timetable of prefrontal cortex white matter (WM) growth and temporal variation in supragranular-enriched gene expression are both used to test whether prefrontal cortex projection development is protracted in humans relative to the timing of other biological pathways. *(a)* A T1 structural MR scan of a chimpanzee brain illustrate boundaries used to measure prefrontal cortex white matter volumes. *(b)* The prefrontal cortex white matter ceases to grow at approximately 1.75 years of age in humans and at approximately 6.7 years of age in chimpanzees In chimpanzees, the prefrontal cortex white matter reaches 60, 70, 80, 90, and 100% of its adult size later than the timing other biological pathways, some of which fall outside the 95% confidence intervals generated from a linear model from these data. *(d-h)* Some supragranular-enriched genes steadily increase in their expression over the course of postnatal development (e., *NEFH, VAMP1*) whereas other genes remain relatively invariant in their expression (e.g., *CRYM*). Chimpanzee age is mapped onto human age to test for deviations in the temporal trajectories in gene expression across species. There were no noticeable differences in temporal trajectories in the expression of these genes between humans and chimpanzees. I compared the coefficient of variation *(i-j)* in the expression of these genes at select age ranges to test for differences in maturational profiles of supragranular-enriched genes [20]. The coefficient of variation in the expression of supragranular-enriched genes was not significantly different between humans and chimpanzees, regardless of whether all age ranges were considered or only late age ranges (10 to 50 years of age) were considered. Overall, these findings converge on the demonstration that prefrontal cortex pathway maturation is not unusually extended in humans.

### Temporal variation in NEFH expression as a marker of white matter pathway maturation

The expression of select supragranular-enriched genes (e.g., *NEFH*) steadily increase at postnatal ages. To ensure the accuracy of these trends and to determine which cell types might account for temporal variation in expression from bulk samples, I quantified temporal changes in *NEFH* expression across layers of the human frontal cortex from in situ hybridization images made available by the Allen Brain Atlas. *NEFH* is of particular interest because it encodes large filaments, is expressed by large neurons, and has been used as a marker of long-range projections (Figs. S5; 26). *NEFH* expression correlates with white matter fraction [Figs. S5-S6, 48]. I randomly selected sites with a grid from downloaded images. I randomly selected frames (1,000 µm thick) aligned along the cortical surface, and I measured *NEFH* expression intensity across layers II–IV versus layers V–VI in the frontal cortex at successive ages, consistent with the methods used in previous work [19,42; Table S4]. I selected 4 sites per individual and quantified *NEFH* expression intensity across layers II–IV and layers V–VI [20,26]. The height of the frames varied with layer thickness. I used cytoarchitecture from adjacent Nissl-stained sections to define layers. I compared *NEFH* expression versus background expression in layers II-IV and V-VI. I then computed a fraction of these two measurements to track temporal changes in the relative expression of *NEFH* across layers (Fig. S6).

## Results

Temporal variation in transcription, anatomy, and behavior find corresponding time points across the lifespan in humans and chimpanzees (Figs. 1, 2). Mapping age with time series data enabled the identification of corresponding time points from gradual changes that span the lifespan (Fig. 2). The use of different methods and scales (e.g., transcription, neuroanatomy, and behavior) yield concordance in corresponding ages between humans and chimpanzees (Fig. 2). Temporal variation in transcription fell within the 95% confidence intervals of other time points in humans versus chimpanzees. The linear model fitting the natural-log-transformed values of age expressed in days after conception in humans and chimpanzees accounted for a large percentage of the variance (1.035x-0.08; R^2^=0.945; p<0.01) and extends beyond 55 year of age in chimpanzees (Fig. 1*c*) and in humans (Fig. 1*d*). The observation that this linear model accounts for 94.5% of the variance demonstrates strong agreements in different scales used to in find corresponding ages in humans and chimpanzees. Chimpanzees start progressing through studied milestones later than humans but progress through these milestones more slowly than in humans. During prenatal development, corresponding time points *occur earlier* in humans than in chimpanzees. As development progresses, the situation is reversed in that corresponding time points actually *occur later* in humans than in chimpanzees (Fig. 5). Together, these data demonstrate the utility of using multiple scales of study to capture corresponding time points across a broad range of ages.

**Fig. 5.**
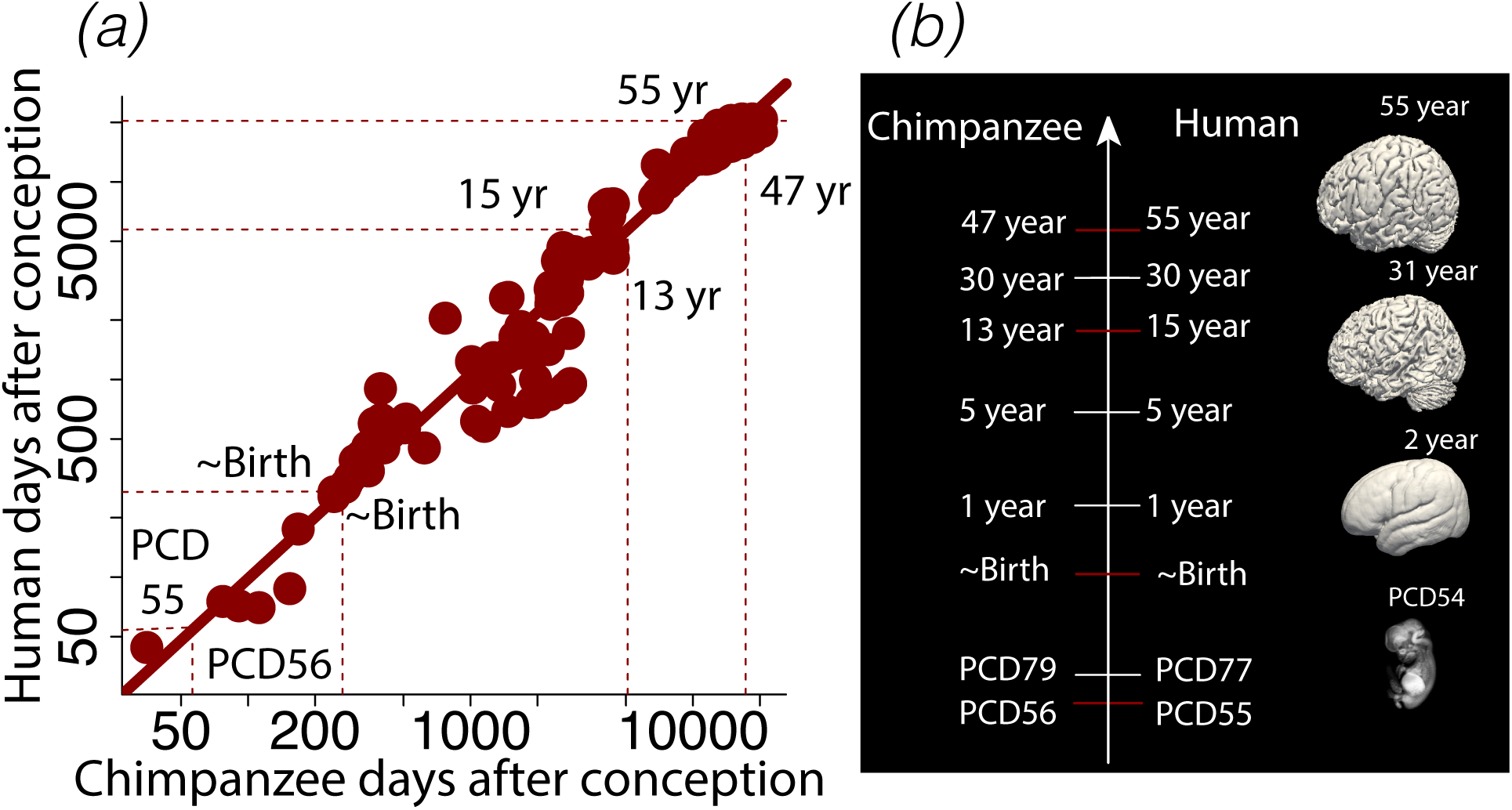
*(a)* Corresponding time points from fetal to postnatal ages across humans and chimpanzees are obtained by regressing the natural-logged time points in days after conception in humans versus chimpanzees. (b) Early in development, humans progress through corresponding time points slightly earlier than in chimpanzees. As humans and chimpanzees start to age, corresponding time points *occur later* in humans than in chimpanzees. Surfaces were extracted from human brain scans at multiple age of development.

### Modifications to the lifespan in primates

I fit a general linear model through the log-transformed time points versus the event scale derived across humans and non-human primates to trace the evolution of developmental schedules across primate species (Fig. 1). The model accounts for a significant percentage of the variance (y=1.62x+2.64; R^2^=98.64; p<0.01) with species and an interaction between species and an event scale as factors. Humans take longer to proceed through transformations than any of the examined primate species (Fig. 1*b*). The schedule of development and aging in chimpanzees resemble humans more than that of macaques and marmosets.

### Evolution of cortical connections

The white matter houses long-range projecting neurons. Comparing timetables of white matter development permits testing for deviations in the timetable of select pathways across species (Fig. 3). White matter prefrontal cortex as well as other pathways (e.g., corpus callosum) growth and cortical regions are protracted in chimpanzees relative to the timing of other biological processes (Figs. 3-4; Fig. S2-S4). The prefrontal cortex white matter ceases to grow at around 6.7 years of age in chimpanzees and 1.75 years in humans. Identified corresponding time points are robust to variation in sample size (Fig. S2*a-d*) and statistical tests (Fig. S2*g-h*) with identified age in the cessation of growth ranging from ∼6 to 8 years in chimpanzees and ∼2 years of age in humans (R^2^∼70-90%; Fig. S2). There is no evidence that prefrontal cortex white matter pathway is protracted in humans relative to the timing of other developmental processes.

### Evolution of transcriptional variation to track long-range projection markers

I considered temporal trajectories in supragranular-enriched gene expression as a means to identify whether the development of frontal cortex circuitry is protracted in humans relative to the timing of other biological pathways. This approach provides an additional means to identify possible species differences in the timeline of the prefrontal cortex pathway maturation between humans and chimpanzees. The expression of some of these supragranular-enriched genes (e.g., *NEFH* and *VAMP1*) steadily increases postnatally in both humans and chimpanzees (Fig. 4*d-k*) whereas other remain relatively invariant. There is no obvious difference in the temporal trajectories in the expression of supragranular-enriched genes across the two species. To detect whether the temporal profiles of supragranular-enriched genes are significantly different in humans compared to chimpanzees, I considered the coefficient of variation across multiple age ranges in humans versus chimpanzees (Fig. 4*i*). I first fit a smooth spline through the log10 of FPKM versus the log10 of age expressed in days after conception for each gene and for each species. A t-test on the coefficient of variation of supragranular-enriched genes between humans and chimpanzees is not significantly different whether all age ranges (t=-0.784; p=0.44, n=13) or 10 to 50 years of age are considered (t=0.101; n=13; p=0.92, n=13, Fig. 4*i*). Comparative analyses of temporal variation in transcription and growth trajectories of the white matter pathway converge on the demonstration that there no evidence of protracted prefrontal cortex projections in humans.

## Discussion

The study employed a multi-scale approach to determine corresponding ages across the lifespan in humans and chimpanzees, which will serve as a resource with which to identify modified developmental programs in the human lineage. Capturing corresponding time points from temporal variation in transcription, anatomy, and behavior ensures accuracy of identified ages across and across an age range that spans the lifespan across species. The synchronism of corresponding ages from different scales is evident given the data’s fit to model accounts for a high percentage (∼94%) of the variance. These data show that, contrary to what has been suggested, the timetable of prefrontal cortex projection maturation is not extended in humans.

### Translating time across the lifespan in humans to chimpanzees

Capturing the timing of development across a broad range of species has identified corresponding ages across prenatal development up to 2 years of age in humans and their equivalent in other species [8, 14,16–18]. This line of work revealed how developmental schedules vary across species and which developmental programs are desynchronized relative to others in select lineages [8, 14,16,19,20,49]. The present study moves the field forward by capturing corresponding ages from gradual changes in brain structure and transcription across the lifespan. Capturing time points from such a broad age ranges shows that the clock setting the pace of aging follows that which sets the pace of development in humans and chimpanzees [14,16–18]. The independent evolution of aging and developmental clocks may become apparent when capturing corresponding time points across a broader range of mammalian species than that used here.

### Evolutionary modifications to developmental programs

Collecting developmental transformations over the course of development identifies modifications to developmental programs in the lineage leading to the emergence of the human brain. One notable example to emerge from this work is the duration of cortical neurogenesis, which is unusually extended in primates relative to other studied mammals (e.g., rodents). Yet, another example is the acceleration of synaptogenesis in humans relative to macaques [8]. The extension in the duration of cortical neurogenesis leads to an amplification of superficial layer neuron numbers and concomitant modifications to cortical circuits [14,16–19]. Another example to emerge from these broad comparative analyses is that birth varies extensively relative to the timing of neural transformations. The use of a single transformation (e.g., birth, puberty, etc.) is not an accurate means to control for variation in development schedules across species but the use of birth is particularly misleading because it has evolved independently of the timetable of neurological programs [22]. The identification of modified developmental programs requires capturing time points from a broad range of developmental programs and across a broad age range to infer modifications to biological programs in the human lineage.

I tested whether developmental programs, such as prefrontal cortex maturation, are protracted in humans compared to chimpanzees. It has been claimed that human prefrontal cortex develops for an unusually long time compared with chimpanzees [2,51,52]. I found no evidence that the prefrontal cortex white matter growth or temporal trajectories in the expression of genes linked to long-range projections (i.e., supragranular-enriched genes) are extended in humans relative to chimpanzees. Rather, cortical pathway maturation is accelerated in humans relative to chimpanzees consistent with previous work, which harnessed temporal variation in transcription to reveal modifications to the timing of developmental programs in humans and non-human primates [8]. These findings exemplify the necessity to control for variation in developmental schedules across species in order to identify modifications to developmental programs within the human lineage [8, 14]. Although the study highlights conservation in the timeline of many biological programs across humans and chimpanzees, not all developmental transformations are conserved in humans versus chimpanzees. Notably, weaning is accelerated in humans [22,53,54]. Life history traits vary relative to the timing of other biological programs and should not be used as a baseline to compare brain development across species.

### Conclusions

This is the first study to demonstrate that a multi-scale approach identifies corresponding ages across the lifespan in human and non-human primates. This novel approach reveals no evidence for protracted prefrontal cortex maturation in humans. Methods other than the ones used here may reveal modification to the timeline of circuitry maturation in the human lineage. Regardless, this study moves us forward by providing a baseline with which to compare ages across humans and chimpanzees across the lifespan, which will enhance our ability to detect modified biological programs in the human lineage.

## Acknowledgements

I thank Melissa Harrington for her support at Delaware State University. Nissl-stained and NEFH+ sections of chimpanzee brains are from the National Chimpanzee Brain Resource (supported by NIH grant NS092988). I used structural MR scans of human fetuses thanks to Dr. Brad Smith at the University of Michigan who provided extended access to prenatal MRI scans. These data are available at http://embryo.soad.umich.edu. Imaging was performed at the Center for In-Vivo Microscopy, Duke University, which was funded by NIH N01-HD-6-3257 P/G F003637.

## Funding

This work was supported by National Institute of General Medical Sciences (NIGMS) grant (5P20GM103653), and a DE-INBRE grant (P20GM103446) to [C.J.C]. The opinions in this article are not necessarily those of the NIH.

## Competing interests

I have no conflict of interest.

## Authors’ contributions

CJC acquired funding, designed the study, analyzed the data, and wrote the manuscript.

---

**Fig. S1.**
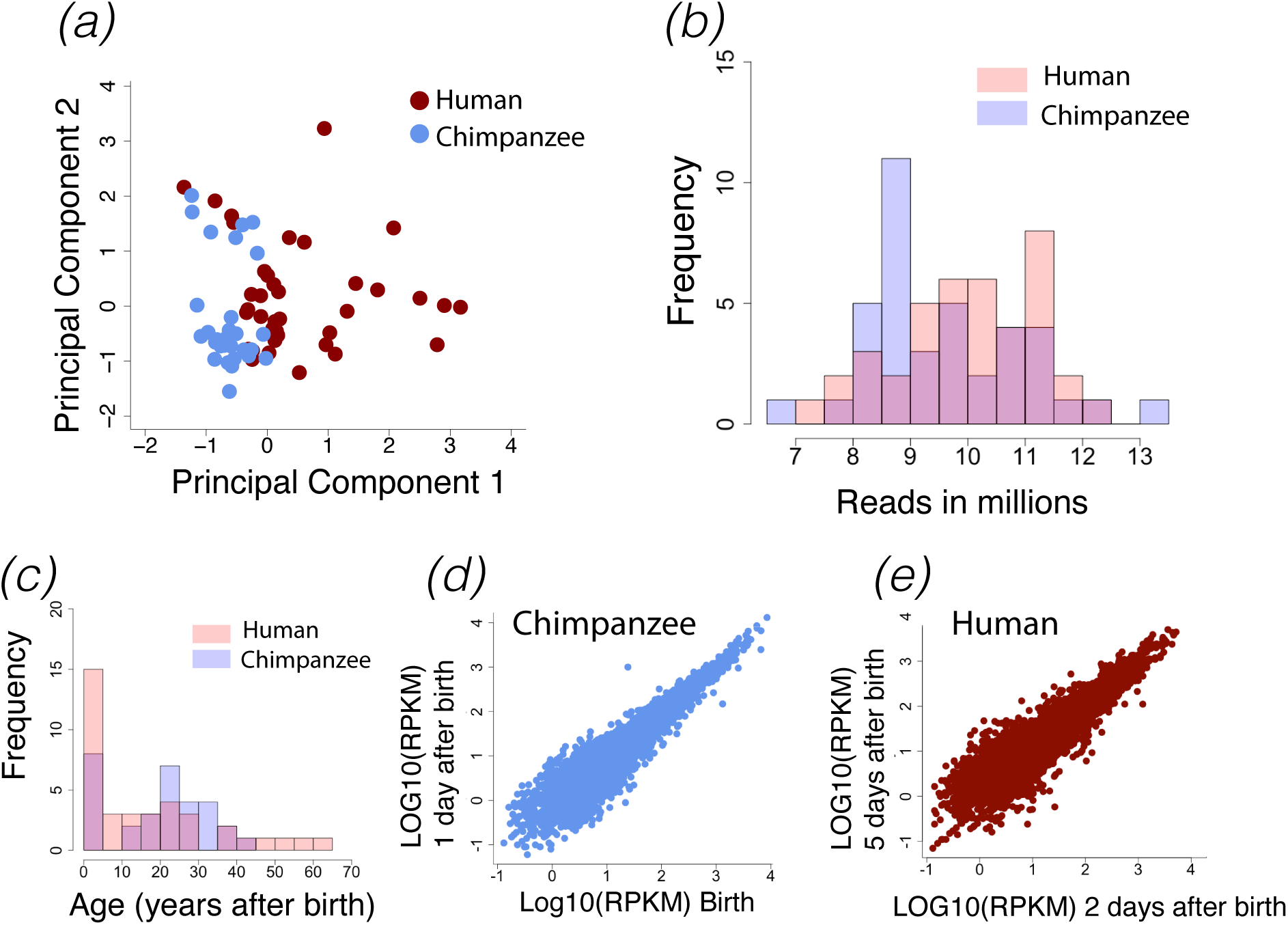
Example of RNA sequencing replicates through the frontal cortex of humans and chimpanzees at different postnatal ages. Minimum expression of 0.5 RPKM was selected across replicates for this figure. (*a*) A principal component analysis on normalized gene expression in humans and chimpanzees shows no obvious outliers. Only expressed genes were selected, and no outliers were removed from the analyses. (*b*) Histogram of mapped RNA sequencing reads and age ranges (*c*) show overlap across species. Biological replicates in chimpanzees and humans are shown in (*d-e*).

**Fig. S2.**
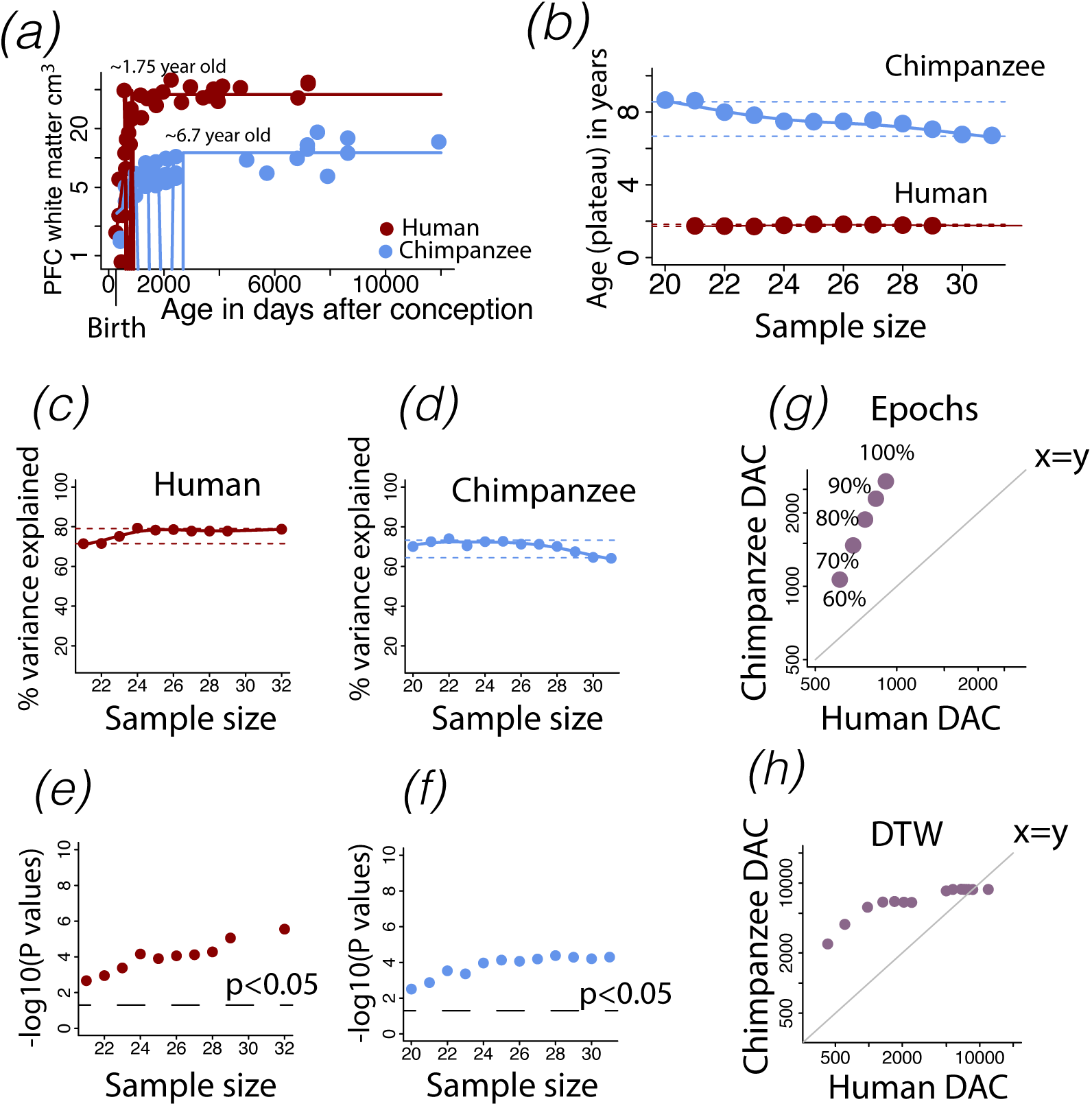
(*a*) I applied a non-linear regression to log-transformed values of the prefrontal cortex white matter volumes versus age expressed in days after conception in humans and chimpanzees. (*a*) I extract when the prefrontal cortex reaches 60%, 70%, 80%, 90%, and 100% of adult volumes, which show that prefrontal cortex growth ceases to grow later in chimpanzees than in humans. (*b-f*) I subsample these data to ensure that corresponding time points extracted from these regressions are not driven by outliers and test for heterochrony using different statistical procedures (g-h). Sample size versus age of identified plateau (b), the percentage of variance accounted for by the model (*c-d*), and test statistics (*e-f*) demonstrate that white matter growth ceases to grow later in chimpanzees that in humans regardless of sample size used (*b*). Time points extracted from either epochs (*g*) or dynamic time warping (*h*) converge to demonstrate that white matter growth occur later in chimpanzees than in humans. Time points about x=y indicate that the events occur later in chimpanzees whereas time points below the regression would indicate that time points occur later in humans than in chimpanzees. Therefore, there is no evidence that prefrontal cortex white matter growth is protracted in humans. DAC=days after conception.

**Fig. S3.**
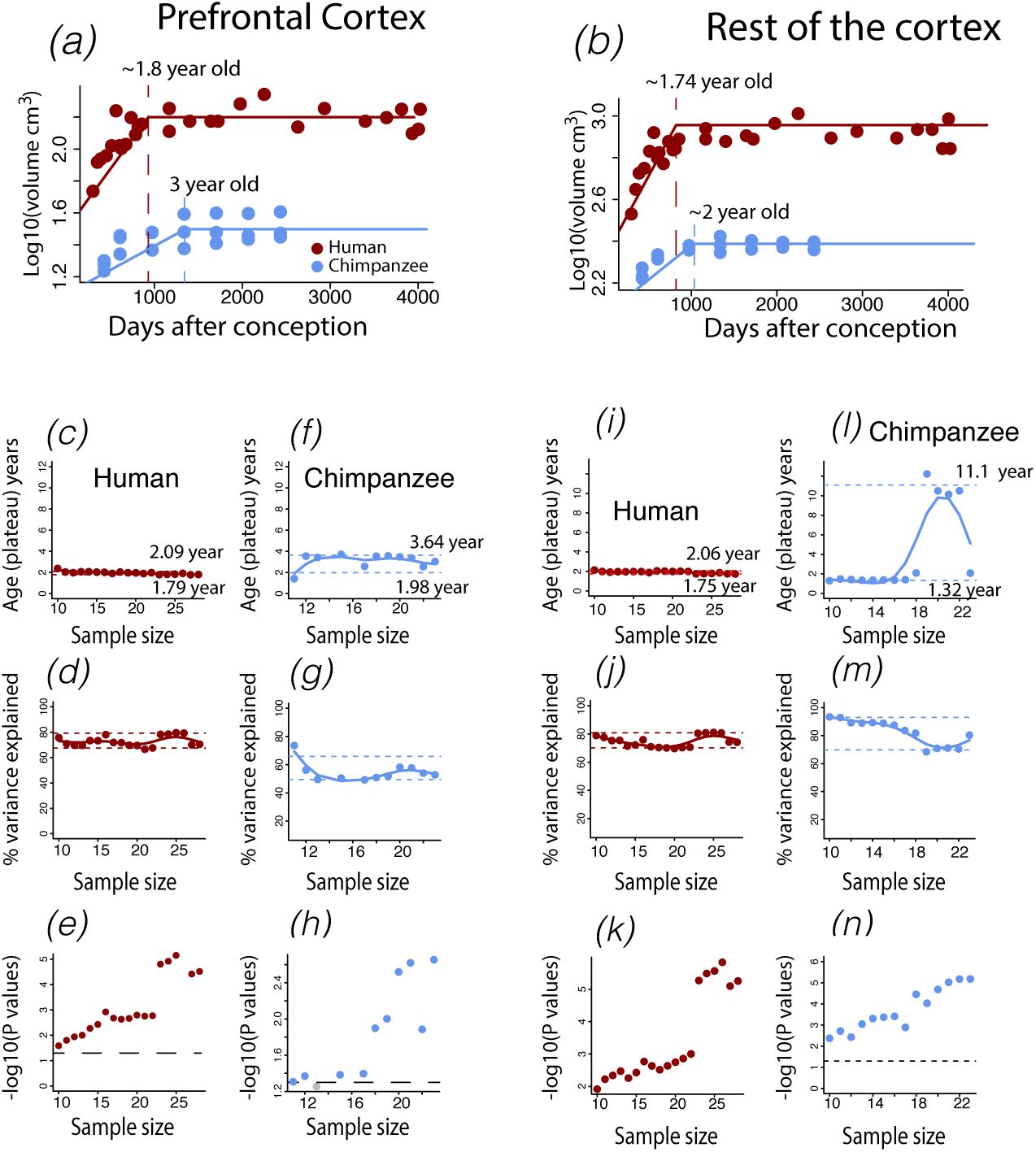
I test how sample size impacts the age of prefrontal cortex (*a*) and rest of the cortex (*b*) growth cessation in humans and in chimpanzees. According to these data, the prefrontal (*a*) and remaining frontal cortex (*b*) cease to grow at around 2 years of age in humans and at approximately 2 to 3 years of age in chimpanzees as shown with vertical lines. I subsample the number of individuals from the non-linear regressions on the log-transformed volumes versus age expressed in days after conception. I extract the age of growth cessation, percentage of variance explained from the model, and test statistics for the human (*c-e*) and chimpanzee prefrontal cortex (*f-h*). I apply the same procedure for the remaining cortex (*i-n*). The 95% confidence intervals extracted for the age of growth cessation and percentage of variance accounted for by these models are shown with horizontal dashed lines. These data show that the remaining frontal cortex ceases to grow around 1.75 to 2 years of age in humans (*c, i*) and around 1.3 to 11 years in chimpanzees (*f, l*). Regardless of sample size, these models account for a significant percentage of the variance in humans (*d-e, j-k*) as in chimpanzees (*g-h, m-n*). There is no evidence that the human prefrontal and remaining frontal cortex is protracted in humans relative to chimpanzees. Contrary to what might be expected, the growth of the prefrontal cortex is protracted in chimpanzees relative to humans.

**Fig. S4.**
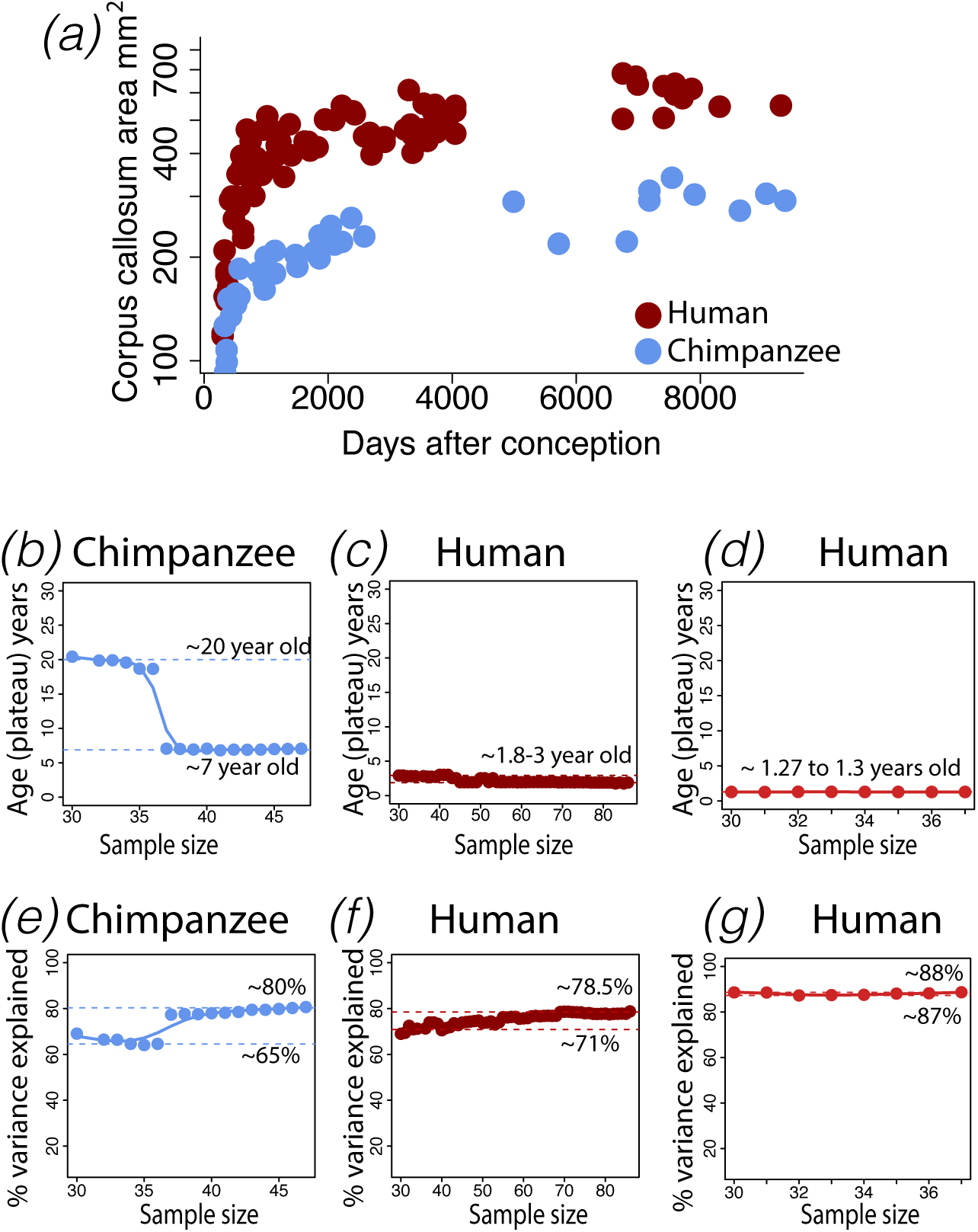
I test how sample size and different data-sets impact the age of identified corpus callosum growth (*a*) cessation in humans and chimpanzees. For each data-set, I subsample the number of individuals and test for a linear plateau applied to the natural-logged values of the corpus callosum area against age expressed in days after conception. I also compare two data-sets to capture when the corpus callosum growth ceases to grow in humans [35, 36, 40]. I test whether sample size impacts estimated age of corpus callosum growth cessation (*b-d*) and percentage of variance explained by the model (e-g). All tested models resulted in significant associations with p<0.05. The age of corpus callosum growth cessation varies extensively in chimpanzees. The 95% confidence intervals (CI) extracted from these data range from 7 to 20 years of age. of corpus callosum cessation growth in humans ranges between ∼1.3 to 3 years of age (*c-d*). Collectively, these data show that there is no evidence that the maturation of this cortical association pathway is protracted in humans relative to chimpanzees.

**Fig. S5.**
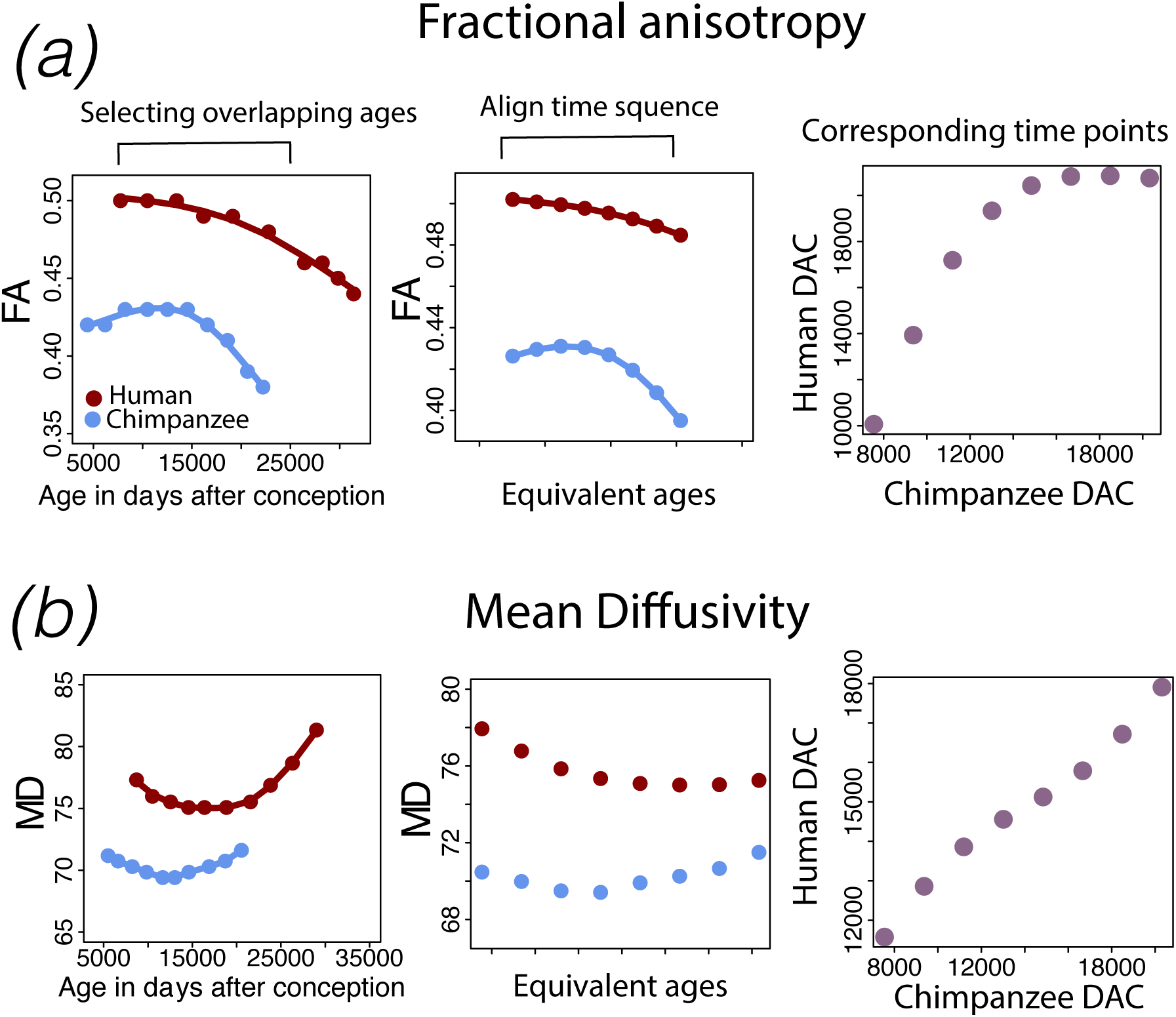
Dynamic time warping applied to temporal variation in cortical fractional anisotropy (FA; *a*) and medial diffusivity (MD; *b*) in humans and chimpanzees was used to capture corresponding time points between humans and chimpanzees. Overlapping age ranges are first selected and these time series data are warped until an optimal alignment is found across humans and chimpanzees, which serves to find corresponding time points between humans and chimpanzees. DAC: days after conception.

**Fig. S6.**
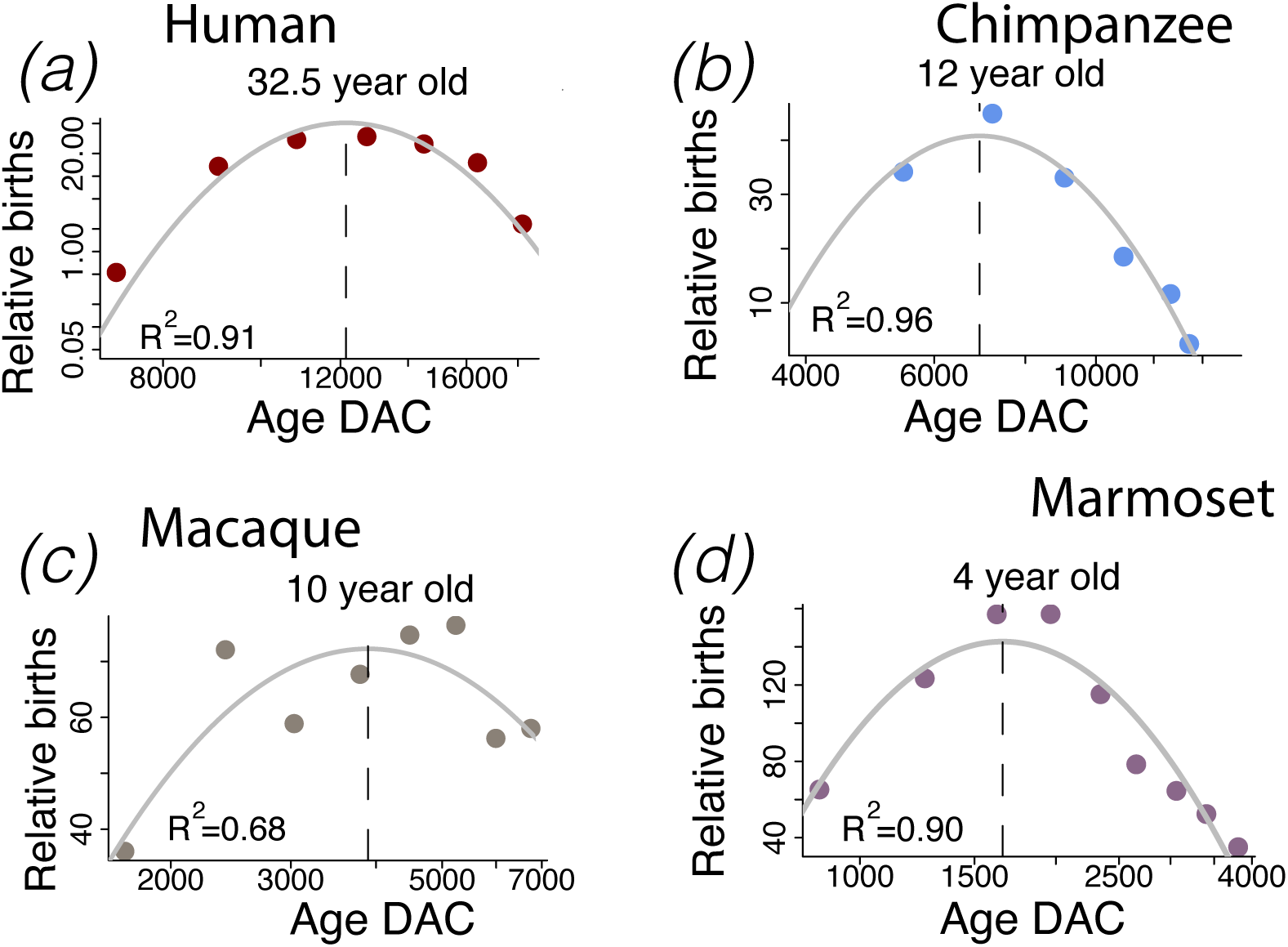
Example of life history time points extracted across humans and non-human primates. I calculated when peaks in relative number of births occur in humans (*a*) chimpanzees (*b*), macaques (*c*), and marmosets (*d*) as a corresponding time point across species.

**Fig. S7.**
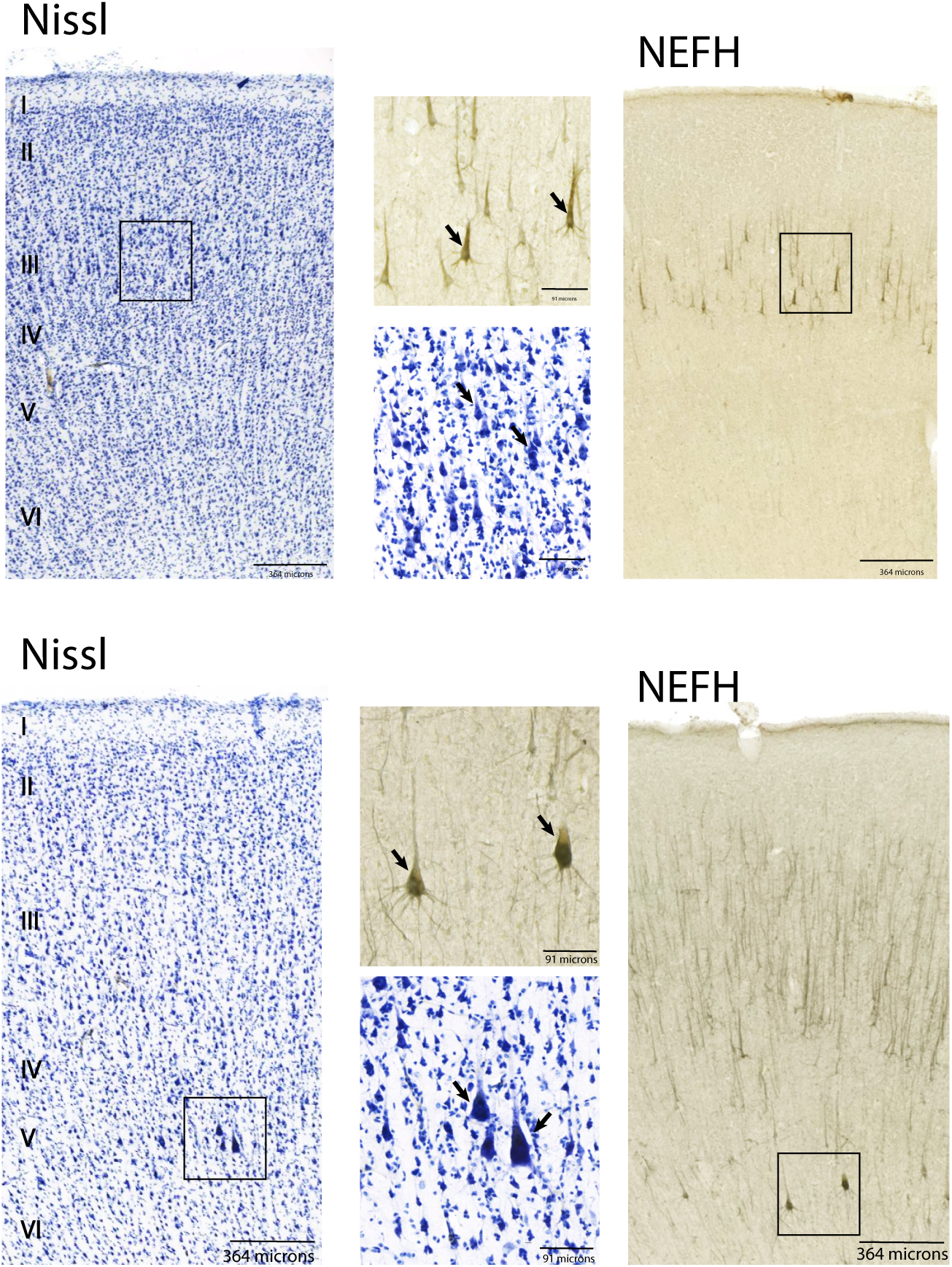
Nissl-stained and NEFH labelled sections from frontal cortical areas in chimpanzees. Across the frontal cortex, cells expressing the NEFH protein are observed in layers III as well as in layer V. These NEFH+ cells are expressed by large neurons in layers III and V. Squares highlight the approximate location of close-up views from low magnification sections. These data are from the National Chimpanzee Brain Resource.

**Fig. S8.**
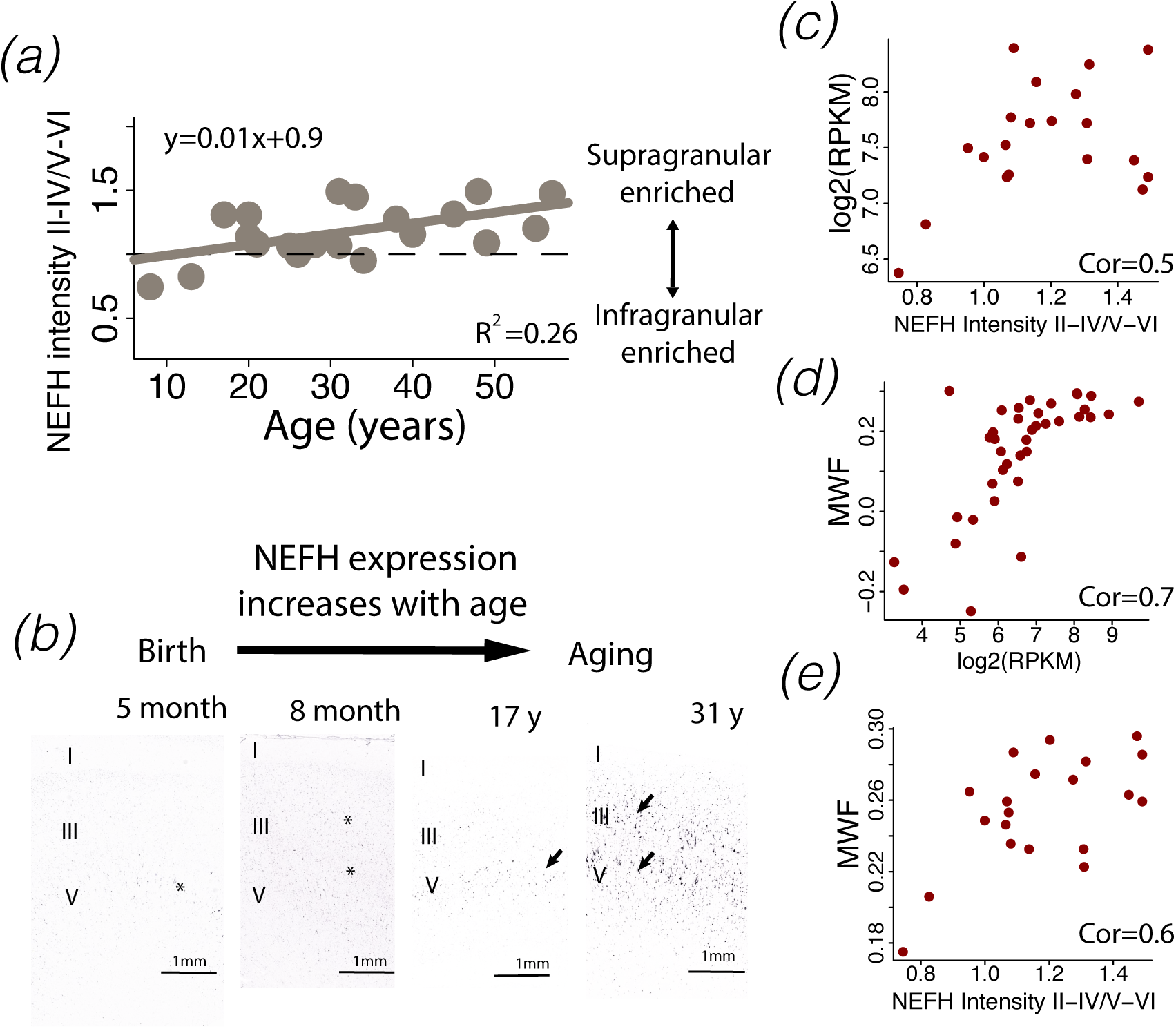
*NEFH* expression steadily increases over postnatal ages in the human frontal cortex (*a-b*). (*c*) Much of the increase in *NEFH* expression from bulk samples is accounted for by a relative increase in *NEFH* expression within supragranular layers. Temporal variation in *NEFH* expression can also be used to track the maturation of white matter pathways through the human frontal cortex. The white matter fraction from the frontal cortex correlates with NEFH expression from bulk samples (*d*) and the relative increase in *NEFH* expression across layers (*e*). Collectively, these data show that protracted developmental trajectories in *NEFH* expression from bulk expression aligns with observations from in situ hybridization and developmental changes in white matter fraction (WMF).

## Notes

### Competing Interest Statement

The authors have declared no competing interest.

## References

1 Somel M, Franz H, Yan Z, Lorenc A, Guo S, Giger T, Kelso J, Nickel B, Dannemann M, Bahn S, et al. 2009 Transcriptional neoteny in the human brain. Proc. Natl. Acad. Sci. U. S. A. 106, 5743–5748. (doi:10.1073/pnas.0900544106)

2 Sakai T, Mikami A, Tomonaga M, Matsui M, Suzuki J, Hamada Y, Tanaka M, Miyabe-Nishiwaki T, Makishima H, Nakatsukasa M, et al. 2011 Differential prefrontal white matter development in chimpanzees and humans. Curr. Biol. 21, 1397–1402. (doi:10.1016/j.cub.2011.07.019)

3 Bufill E, Agustí J, Blesa R. 2011 Human neoteny revisited: the case of synaptic plasticity. Am. J. Hum. Biol. 23, 729–739. (doi:10.1002/ajhb.21225)

4 Miller DJ, Duka T, Stimpson CD, Schapiro SJ, Baze WB, McArthur MJ, Fobbs AJ, Sousa AMM, Sestan N, Wildman DE, et al. 2012 Prolonged myelination in human neocortical evolution. Proc. Natl. Acad. Sci. U. S. A. 109, 16480–16485. (doi:10.1073/pnas.1117943109)

5 Liu X, Somel M, Tang L, Yan Z, Jiang X, Guo S, Yuan Y, He L, Oleksiak A, Zhang Y, et al. 2012 Extension of cortical synaptic development distinguishes humans from chimpanzees and macaques. Genome Res. 22, 611–622. (doi:10.1101/gr.127324.111)

6 Sakai T, Matsui M, Mikami A, Malkova L, Hamada Y, Tomonaga M, Suzuki J, Tanaka M, Miyabe-Nishiwaki T, Makishima H, et al. 2013 Developmental patterns of chimpanzee cerebral tissues provide important clues for understanding the remarkable enlargement of the human brain. Proc. Biol. Sci. 280, 20122398. (doi:10.1098/rspb.2012.2398)

7 Bianchi S, Stimpson CD, Duka T, Larsen MD, Janssen WGM, Collins Z, Bauernfeind AL, Schapiro SJ, Baze WB, McArthur MJ, et al. 2013 Synaptogenesis and development of pyramidal neuron dendritic morphology in the chimpanzee neocortex resembles humans. Proc. Natl. Acad. Sci. U. S. A. 110 Suppl 2, 10395–10401. (doi:10.1073/pnas.1301224110)

8 Zhu Y, Sousa AMM, Gao T, et al. 2018. Spatiotemporal transcriptomic divergence across human and macaque brain development. Science. 362, eaat8077. (doi:10.1126/science.aat8077)

9 Charvet CJ, Finlay BL. 2018 Comparing adult hippocampal neurogenesis across species: Translating time to predict the tempo in humans. Front. Neurosci. 12, 706. (doi:10.3389/fnins.2018.00706)

10 Mora-Bermúdez F, Badsha F, Kanton S, Camp JG, Vernot B, Köhler K, Voigt B, Okita K, Maricic T, He Z, et al. 2016 Differences and similarities between human and chimpanzee neural progenitors during cerebral cortex development. eLife 5, e18683. (doi:10.7554/eLife.18683)

11 Kanton S, Boyle MJ, He Z, Santel M, Weigert A, Sanchís-Calleja F, Guijarro P, Sidow L, Fleck JS, Han D, et al. 2019 Organoid single-cell genomic atlas uncovers human-specific features of brain development. Nature 574, 418–422. (doi:10.1038/s41586-019-1654-9)

12 Kanton S, Treutlein B, Camp JG. 2020 Single-cell genomic analysis of human cerebral organoids. Methods Cell Biol. 159, 229–256. (doi:10.1016/bs.mcb.2020.03.013)

13 Tang H, Tang Y, Zeng T, Chen L. 2020 Gene expression analysis reveals the tipping points during infant brain development for human and chimpanzee. BMC Genomics 21, 74. (doi:10.1186/s12864-020-6465-8).

14 Finlay B, Darlington R. 1995 Linked regularities in the development and evolution of mammalian brains. Science 268, 1578–1584. (doi:10.1126/science.7777856)

15 Clancy B, Darlington RB, Finlay BL. 2001 Translating developmental time across mammalian species. Neuroscience 105, 7–17. (doi:10.1016/s0306-4522(01)00171-3)

16 Clancy B, Finlay BL, Darlington RB, Anand KJS. 2007 Extrapolating brain development from experimental species to humans. Neurotoxicology 28, 931–937. (doi:10.1016/j.neuro.2007.01.014)

17 Workman AD, Charvet CJ, Clancy B, Darlington RB, Finlay BL. 2013 Modeling transformations of neurodevelopmental sequences across mammalian species. J. Neurosci. 33, 7368–7383. (doi:10.1523/JNEUROSCI.5746-12.2013)

18 Charvet CJ, Hof PR, Raghanti MA, Van Der Kouwe AJ, Sherwood CC, Takahashi E. 2017 Combining diffusion magnetic resonance tractography with stereology highlights increased cross-cortical integration in primates. J. Comp. Neurol. 525, 1075–1093. (doi:10.1002/cne.24115)

19 Charvet CJ, Šimić G, Kostović I, Knezović V, Vukšić M, Babić Leko M, Takahashi E, Sherwood CC, Wolfe MD, Finlay BL. 2017 Coevolution in the timing of GABAergic and pyramidal neuron maturation in primates. Proc. Biol. Sci. 284, 20171169. (doi:10.1098/rspb.2017.1169)

20 Hendy JP, Takahashi E, van der Kouwe AJ, Charvet CJ. 2020 Brain wiring and supragranular-enriched genes linked to protracted human frontal cortex development. Cereb. Cortex. In Press. (doi:10.1093/cercor/bhaa135)

21 Pattabiraman K, Muchnik SK, Sestan N. 2020. The evolution of the human brain and disease susceptibility. Curr Opin Genet Dev. 65:91–97. doi:10.1016/j.gde.2020.05.004.

22 Hawkes K, Finlay BL. 2018 Mammalian brain development and our grandmothering life history. Physiol. Behav. 193, 55–68. (doi:10.1016/j.physbeh.2018.01.013)

23 Finlay BL, Hersman MN, Darlington RB. 1998 Patterns of vertebrate neurogenesis and the paths of vertebrate evolution. Brain Behav. Evol. 52, 232–242. (doi:10.1159/000006566)

24 Finlay BL, Huang K. 2020 Developmental duration as an organizer of the evolving mammalian brain: scaling, adaptations, and exceptions. Evol. Dev. 22, 181–195. (doi:10.1111/ede.12329)

25 Zeng H, Shen EH, Hohmann JG, Oh SW, Bernard A, Royall JJ, Glattfelder KJ, Sunkin SM, Morris JA, Guillozet-Bongaarts AL, et al. 2012 Large-scale cellular-resolution gene profiling in human neocortex reveals species-specific molecular signatures. Cell 149, 483–496. (doi:10.1016/j.cell.2012.02.052)

26 Charvet CJ, Palani A, Kabaria P, Takahashi E. 2019 Evolution of brain connections: Integrating diffusion MR tractography with gene expression highlights increased corticocortical projections in primates. Cereb. Cortex 29, 5150–5165. (doi:10.1093/cercor/bhz054)

27 Charvet CJ. 2020 Closing the gap from transcription to the structural connectome enhances the study of connections in the human brain. Dev. Dyn. (doi:10.1002/dvdy.218)

28 Krienen FM, Yeo BTT, Ge T, Buckner RL, Sherwood CC. 2016 Transcriptional profiles of supragranular-enriched genes associate with corticocortical network architecture in the human brain. Proc. Natl. Acad. Sci. U. S. A. 113, E469–E478. (doi:10.1073/pnas.1510903113)

29 Chevalier-Skolnikoff S. 1983 Sensorimotor development in orang-utans and other primates. J. Hum. Evol. 12, 545–561. (doi:10.1016/s0047-2484(83)80034-7)

30 Macho GA, Wood BA. 1995 The role of time and timing in hominid dental evolution. Evol. Anthropol.: Issues News Rev. 4, 17–31. (doi:10.1002/evan.1360040105)

31 Caro TM, Sellen DW, Parish A, Frank R, Brown DM, Voland E, Mulder MB. 1995 Termination of reproduction in nonhuman and human female primates. Int. J. Primatol. 16, 205–220. (doi:10.1007/bf02735478)

32 Kurjak A, Pooh RK, Merce LT, Carrera JM, Salihagic-Kadic A, Andonotopo W. 2005 Structural and functional early human development assessed by three-dimensional and four-dimensional sonography. Fertil. Steril. 84, 1285–1299. (doi:10.1016/j.fertnstert.2005.03.084)

33 Mizuno Y, Takeshita H, Matsuzawa T. 2006 Behavior of infant chimpanzees during the night in the first 4 months of life: smiling and suckling in relation to behavioral state. Infancy 9, 221–240. (doi:10.1207/s15327078in0902_7)

34 Lüchinger AB, Hadders-Algra M, van Kan CM, de Vries JIP. 2008 Fetal onset of general movements. Pediatr. Res. 63, 191–195. (doi:10.1203/pdr.0b013e31815ed03e)

35 Garwicz M, Christensson M, Psouni E. 2009 A unifying model for timing of walking onset in humans and other mammals. Proc. Natl. Acad. Sci. U. S. A. 106, 21889–21893. (doi:10.1073/pnas.0905777106)

36 Chen X, Errangi B, Li L, Glasser MF, Westlye LT, Fjell AM, Walhovd KB, Hu X, Herndon JG, Preuss TM, et al. 2013 Brain aging in humans, chimpanzees (Pan troglodytes), and rhesus macaques (Macaca mulatta): magnetic resonance imaging studies of macro- and microstructural changes. Neurobiol. Aging 34, 2248–2260. (doi:10.1016/j.neurobiolaging.2013.03.028)

37 Hikishima K, Sawada K, Murayama AY, Komaki Y, Kawai K, Sato N, Inoue T, Itoh T, Momoshima S, Iriki A, et al. 2013 Atlas of the developing brain of the marmoset monkey constructed using magnetic resonance histology. Neuroscience 230, 102–113. (doi:10.1016/j.neuroscience.2012.09.053)

38 Narayanan DZ. 2015 The organization of action in the fetus Prenatal development of orofacial movements in marmoset monkeys [Doctoral dissertation]. Princeton, NJ: Princeton University.

39 Takeshita H, Hirata S, Sakai T, Myowa-Yamakoshi M. 2016 Fetal behavioral development and brain growth in chimpanzees versus humans: a view from studies with 4d ultrasonography. In Fetal development: research on brain and behavior, environmental influences, and emerging technologies (eds. N. Reissland, BS. Kisilevsky), pp. 67–83. Cham, Switzerland: Springer International Publishing.

40 Ausderau KK, Dammann C, McManus K, Schneider M, Emborg ME, Schultz-Darken N. 2017 Cross-species comparison of behavioral neurodevelopmental milestones in the common marmoset monkey and human child. Dev. Psychobiol. 59, 807–821. (doi:10.1002/dev.21545)

41 Sakai T, Komaki Y, Hata J, Okahara J, Okahara N, Inoue T, Mikami A, Matsui M, Oishi K, Sasaki E, et al. 2017 Elucidation of developmental patterns of marmoset corpus callosum through a comparative MRI in marmosets, chimpanzees, and humans. Neurosci. Res. 122, 25–34. (doi:10.1016/j.neures.2017.04.001)

42 Sakai T, Mikami A, Suzuki J, Miyabe-Nishiwaki T, Matsui M, Tomonaga M, Hamada Y, Matsuzawa T, Okano H, Oishi K. 2017 Developmental trajectory of the corpus callosum from infancy to the juvenile stage: comparative MRI between chimpanzees and humans. PLoS One 12, e0179624. (doi:10.1371/journal.pone.0179624)

43 Lonsdorf EV, Stanton MA, Pusey AE, Murray CM. 2020 Sources of variation in weaned age among wild chimpanzees in Gombe National Park, Tanzania. Am. J. Phys. Anthropol. 171, 419–429. (doi:10.1002/ajpa.23986)

44 Bründl AC, Tkaczynski PJ, Nohon Kohou G, Boesch C, Wittig RM, Crockford C. 2020 Systematic mapping of developmental milestones in wild chimpanzees. Dev. Sci. e12988. (doi:10.1111/desc.12988)

45 Lister R, Mukamel EA, Nery JR, Urich M, Puddifoot CA, Johnson ND, Lucero J, Huang Y, Dwork AJ, Schultz MD, et al. 2013 Global epigenomic reconfiguration during mammalian brain development. Science 341, 1237905. (doi:10.1126/science.1237905)

46 Charvet CJ, Das A, Song JW, Tindal-Burgess DJ, Kabaria P, Dai G, Kane T, Takahashi E. 2020. High Angular Resolution Diffusion MRI Reveals Conserved and Deviant Programs in the Paths that Guide Human Cortical Circuitry. Cereb Cortex 30, 1447–1464. doi:10.1093/cercor/bhz17841.

47 Richardson H, Lisandrelli G, Riobueno-Naylor A, Saxe R. 2018. Development of the social brain from age three to twelve years. Nature communications 9, 1027.

48 Deoni SCL, Dean DC, 3rd, O’Muircheartaigh J, Dirks H, Jerskey BA. 2012 Investigating white matter development in infancy and early childhood using myelin water faction and relaxation time mapping. NeuroImage 63, 1038–1053. (doi:10.1016/j.neuroimage.2012.07.037)

49 Hof PR, Nimchinsky EA, Morrison JH. 1995 Neurochemical phenotype of corticocortical connections in the macaque monkey: quantitative analysis of a subset of neurofilament protein-immunoreactive projection neurons in frontal, parietal, temporal, and cingulate cortices. J. Comp. Neurol. 362, 109–133. (doi:10.1002/cne.903620107)

50 Nguyen MQ, Wu Y, Bonilla LS, von Buchholtz LJ, Ryba NJP. 2017 Diversity amongst trigeminal neurons revealed by high throughput single cell sequencing. PLoS One 12, e0185543. (doi:10.1371/journal.pone.0185543)

51 Finlay BL. 2019 Human exceptionalism, our ordinary cortex and our research futures. Dev Psychobiol. 61, 317–322. doi:10.1002/dev.21838

52 Barton RA, Venditti C. 2013 Human frontal lobes are not relatively large. Proc. Natl. Acad. Sci. U. S. A. 110, 9001–9006. (doi:10.1073/pnas.1215723110)

53 Galdikas BMF, Wood JW. 1990 Birth spacing patterns in humans and apes. Am. J. Phys. Anthropol. 83, 185–191. (doi:10.1002/ajpa.1330830207)

54 Hawkes K, O’Connell JF, Jones NG, Alvarez H, Charnov EL. 1998 Grandmothering, menopause, and the evolution of human life histories. Proc. Natl. Acad. Sci. U. S. A. 95, 1336–1339. (doi:10.1073/pnas.95.3.1336)

